# Structural basis for DARC binding in reticulocyte invasion by *Plasmodium vivax*

**DOI:** 10.1101/2023.02.28.530453

**Authors:** Re’em Moskovitz, Tossapol Pholcharee, Sophia M. DonVito, Bora Guloglu, Edward Lowe, Franziska Mohring, Robert W. Moon, Matthew K. Higgins

**Affiliations:** Department of Biochemistry, University of Oxford, South Parks Road, Oxford, OX1 3QU, UK; London School of Hygiene and Tropical Medicine, Keppel Street, London, WC1E 7HT, UK

## Abstract

The symptoms of malaria occur during the blood stage of infection, when the parasite replicates within human red blood cells. The human malaria parasite, *Plasmodium vivax*, selectively invades reticulocytes in a process which requires an interaction between the ectodomain of the human DARC receptor and the *Plasmodium vivax* Duffy-binding protein, PvDBP. Previous studies have revealed that a small helical peptide from DARC binds to region II of PvDBP (PvDBP-RII). However, it is also known that sulphation of tyrosine residues on DARC affects its binding to PvDBP and these residues were not observed in previous structures. We have therefore determined the structure of PvDBP-RII bound to sulphated DARC peptide, showing that a sulphate on tyrosine 41 binds to a charged pocket on PvDBP-RII. We use molecular dynamics simulations, affinity measurements and growth-inhibition experiments in parasites to confirm the importance of this interaction. We also reveal the epitope for vaccine-elicited growth-inhibitory antibody, DB1. This provides a complete understanding of the binding of PvDBP-RII to DARC and will guide the design of vaccines and therapeutics to target this essential interaction.

## Introduction

In many regions of the world, *Plasmodium vivax* is the predominant parasite that causes human malaria. Leading to around 14.5 million diagnosed cases each year, it accounts for 75% of malaria in the Americas and 50% in South East Asia^1^. With studies suggesting that the impact of this parasite is under-estimated^2,3^, morbidity caused by *Plasmodium vivax* is a major global health problem and an effective vaccine is urgently required to contribute to the malaria eradication effort.

The symptoms of malaria occur when parasites replicate within the blood. In the case of *Plasmodium vivax*, invasion of red blood cells is highly dependent on the reticulocyte Duffy antigen/receptor for chemokines (DARC). The importance of DARC as an invasion receptor^4^ was first shown over 45 years ago, with the demonstration that the related parasite, *Plasmodium knowlesi*, is unable to invade blood cells taken from individuals with the Duffy-negative phenotype^*5*^. These people have a mutation in their DARC gene which makes the receptor undetectable on circulating reticulocytes. Duffy negativity was also found to prevent African-Americans from being infected with *Plasmodium vivax* through mosquito bite^6^. The widespread prevalence of *Plasmodium vivax* across the world, together with its much-reduced occurrence in regions of Africa where Duffy-negativity is widespread, highlight the importance of DARC as a determinant of *Plasmodium vivax* susceptibility^7^. Indeed, while *Plasmodium vivax* is occasionally capable of infecting Duffy-negative individuals, this is linked to lower parasite load and more than 15-fold reduced likelihood of causing clinical disease^8-10^. Preventing DARC-mediated invasion remains a primary approach to prevent clinical vivax malaria.

The parasite receptor for DARC is the Duffy binding protein, PvDBP^11^ and multiple lines of evidence highlight the important nature of their interaction. In the closely related, genetically tractable parasite, *Plasmodium knowlesi*, knockout of PkDBPα prevents invasion of Duffy-positive erythrocytes *in vitro*^12-14^. While PvDBP knockout in *Plasmodium vivax* has not been technically possible, due to challenges with maintaining this parasite in culture conditions, immunisation of mice, rabbits and non-human primates with the RII region of PvDBP (PvDBP-RII) induces inhibitory antibodies that block binding of PvDBP to DARC^15,16^. In humans, high-titres of naturally-acquired antibodies that target PvDBP-RII and prevent DARC binding *in vitro*, are associated with reduced risk of *Plasmodium vivax* infection^17^, lower parasite densities following invasion and decreased risk of clinical malaria^18,19^. Immunisation of human volunteers with recombinant viral vectors expressing PvDBP-RII induces strain-transcending antibodies which prevent recombinant PvDBP-RII from binding to DARC while human antibodies, derived from either vaccination or natural infection, are inhibitory of invasion^13,20^. PvDBP is therefore the primary blood stage vaccine candidate to prevent vivax malaria.

With both intact PvDBP and DARC proving challenging to produce, molecular studies have used smaller fragments to narrow down functionally important regions^21^, with a ∼350 amino acid residue Duffy-binding-like (DBL) domain known as PvDBP-RII, shown to bind DARC^22^. PvDBP-RII binds to the sixty-residue N-terminal ectodomain of DARC^23^. The only structural insight into this interaction comes from structures of PvDBP-RII bound to an 11-residue helical peptide (DARC_19-30_). However, DARC is post-translationally modified by sulphation of two tyrosine residues, Y30 and Y41 and mutation of Y41 to phenylalanine led to reduced PvDBP binding^24^. These residues are not observed in existing structures, most likely because the DARC ectodomain used was bacterially expressed and therefore lacked tyrosine sulphation^24,25^. Mutations have also been described in regions of PvDBP which do not contact DARC_19-30_ but which are known to contribute to full-length DARC binding^26,27^ and a polymorphism in residue 42 of DARC is the sole change in the Fy^a^ blood group, which reduces the likelihood and severity of *Plasmodium vivax* infection^28^. Indeed, NMR experiments indicate that residues from 14-43 of DARC show chemical shifts on binding to PvDBP-RII^29^. The full interaction between PvDBP and DARC therefore remains to be defined.

Structural approaches have also started to reveal how the PvDBP-DARC interaction can be blocked by mouse and human monoclonal antibodies. Two human monoclonals bind the region of PvDBP involved in DARC_19-30_ binding and dimerisation^20^. In contrast, a broadly inhibitory human monoclonal, from a vaccinated human volunteer, binds away from this site^13^, as do inhibitory mouse antibodies^30^. How do these work without directly blocking the PvDBP-DARC interaction?

In this study, we determine the structural basis for the interaction of PvDBP with the DARC ectodomain and the human invasion blocking antibody, DB1. By using DARC proteins expressed in human cells, we define the binding site for sulphated tyrosine 41. This reveals the molecular basis for the role of tyrosine sulphation in the essential interaction between PvDBP and DARC in reticulocyte invasion by *Plasmodium vivax*.

## Results

### Structural characterisation of the PvDBP-DARC interaction

To determine the structure of PvDBP-RII bound to sulphated DARC ectodomain we started by expressing the N-terminal 60 residues of DARC (DARC_ecto_) in HEK293 cells. This was purified and assessed by mass spectrometry, with the predominant peak having the mass expected for the peptide with the addition of two sulphates. We also expressed PvDBP-RII in HEK293 cells and this was mixed with DARC_ecto_ for crystallisation. As no crystals formed, we attempted to use Fab fragments from monoclonal antibodies which bind to PvDBP-RII, but do not prevent it from binding to DARC_ecto_, as crystallisation chaperones^13^. A complex of PvDBP-RII, DARC_ecto_ and the Fab fragment of the DB1 monoclonal antibody^13^ formed crystals which diffracted to 2.49Å resolution. Molecular replacement using the PvDBP-RII^13^ structure as a search model provided phase information and the resultant electron density map revealed density for residues 19-47 of DARC_ecto_ (Figure 1, Supplementary Table 1).

**Figure 1:**
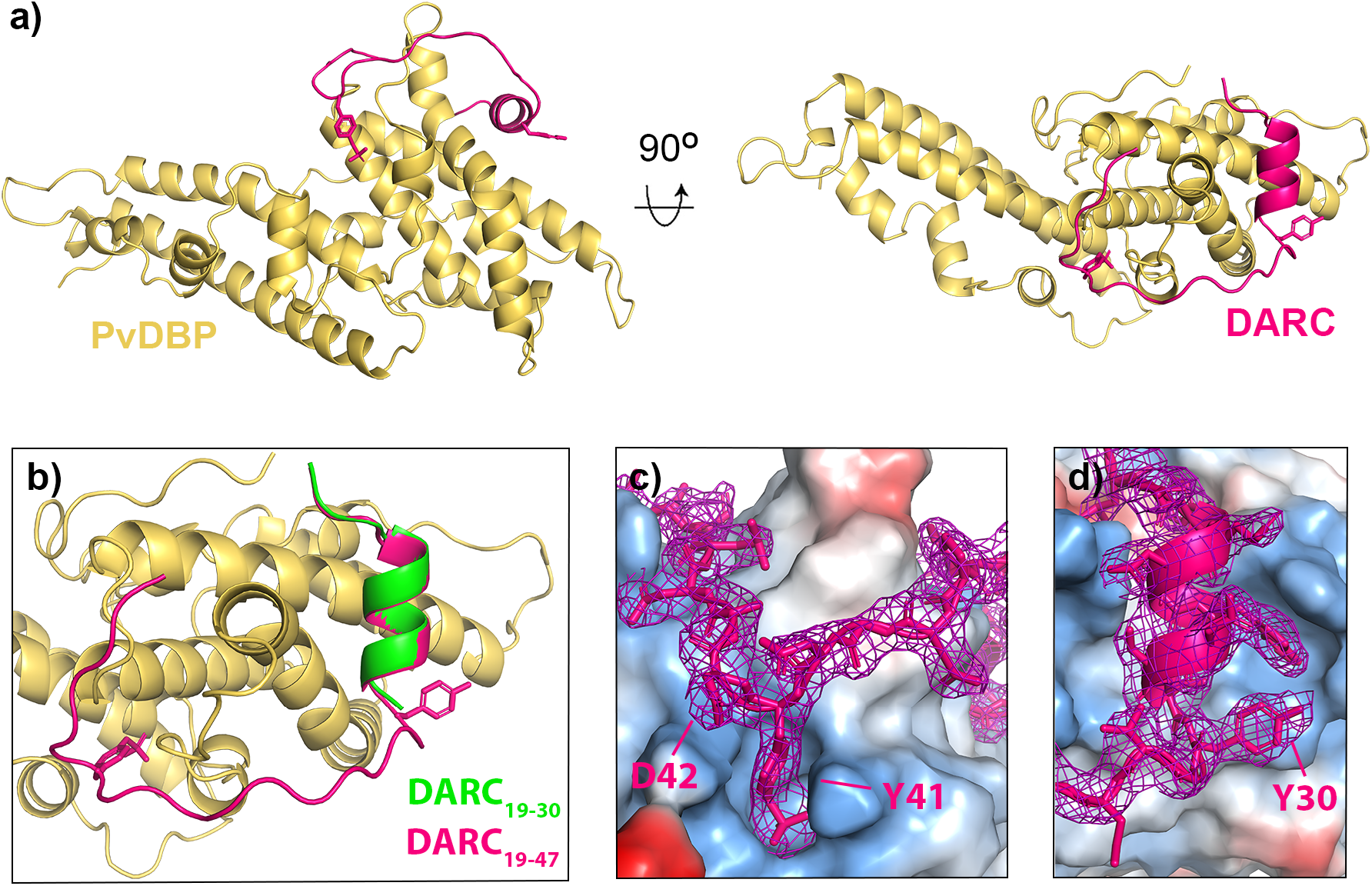
The structure of PvDBP-RII bound to sulphated DARC ectodomain. **a)** The structure of PvDBP-RII (yellow) bound to the DARC ectodomain (pink). **b)** A close up of the DARC ectodomain, showing residues 19-47 of the sulphated ectodomain in pink, overlayed with residues 19-30 of the non-sulphated ectodomain in green^29^. **c)** A close up of residue 41 of DARC, with DARC and the electron density surrounding DARC in pink and PvDBP-RII as a surface coloured by electrostatics (blue as positive charge and red as negative). **d)** A close up of residue 30 of DARC, with DARC and the electron density surrounding DARC in pink and PvDBP-RII as a surface coloured by electrostatics (blue as positive charge and red as negative).

A comparison with the previously determined structure^29^ of PvDBP-RII bound to DARC_ecto_ expressed in bacteria shows DARC residues 19-30 to adopt the same binding mode in both cases, forming an *α*-helix which packs against subdomain 2 of PvDBP-RII (Figure 1b). While this is the only region of DARC observed in the previous study, we now also observe clear density for residues 31-47. These form an elongated peptide which wraps, in a horse-shoe shaped trajectory, around a protrusion on the surface of PvDBP-RII subdomain 2. Clear density was observed for each residue, including Y41 and the sulphate on Y41 (Figure 1c). Also visible is the side chain of residue D42, which is polymorphic (D42G) in the Fy^a^/Fy^b^ blood group variant (Figure 1c). Residues 48-60 are not visible in the crystal structure, suggesting a disordered linker between the PvDBP-RII-binding region of the ectodomain and the DARC transmembrane region, which starts at residue 61.

### Sulphated Y41 binds to a stable binding pocket on PvDBP while Y30 is dynamic

The structure allows us to rationalise the role of sulphation of Y30 and Y41 of DARC on the interaction with PvDBP. A previous study has shown that mutation of Y30 to F, which removes the hydroxyl group, as well as the potential for sulphation, did not affect PvDBP-RII binding^24^. Indeed, in our structure, the Y30 side chain was clearly visible in the electron density, but no evidence was seen for sulphation, suggesting either that the sulphate was not present, or that it was disordered in the electron density (Figure 1d). The side chain did not contact PvDBP-RII, but instead interacts with a neighbouring DB1 Fab fragment through crystal contacts.

In contrast, the side chain of Y41, together with clear density for a sulphate group, was observed. This docks into a positively charged pocket on the PvDBP-RII surface, where the sulphate makes direct salt-bridges with K301 and R304 (Figure 1c). This is consistent with sulphated Y41 (Y41-S) forming an important part of the binding interface. Indeed, the more substantial Y41F mutation, which removes both sulphate and hydroxyl group, reduces PvDBP binding^24^ and this interaction is likely to be important for residues 31-47 to adopt their correct binding conformation.

We next assessed the stability of the binding modes of Y30 and Y41 using metadynamics simulations, allowing us to generate the relative free energy landscapes of sulphated and non-sulphated versions of Y30 and Y41^31,32^. These simulations were run for three different molecular models, generated from our crystal structure. These were sulphated on both Y30 and Y41, sulphated on just Y30 and sulphated on just Y41. In each case, we analysed the ensemble of structures generated during the simulation, the free energy surfaces and the number and duration of contacts formed between each tyrosine residue and PvDBP-RII Figure 2).

**Figure 2:**
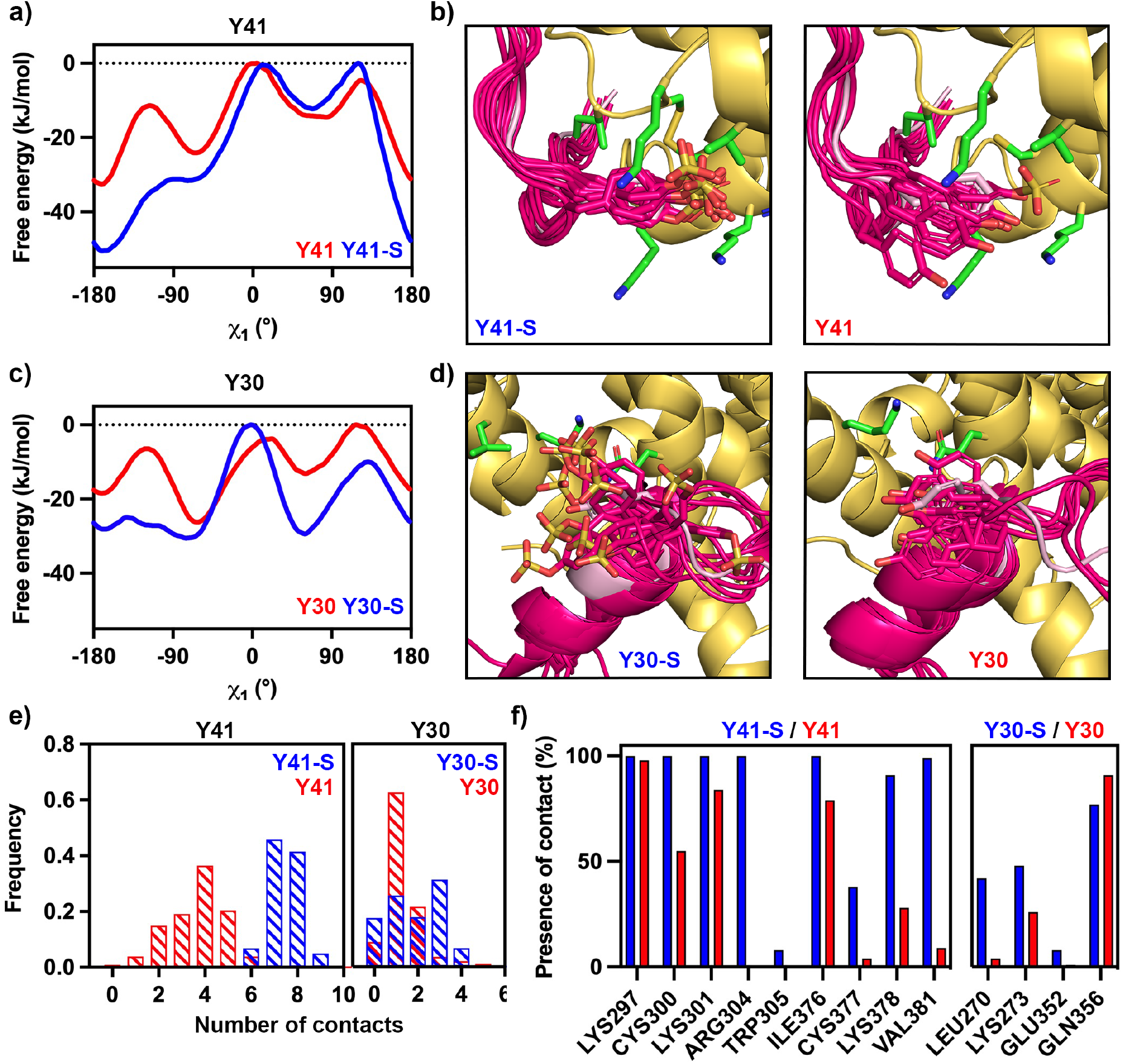
Molecular dynamics simulations indicate ordered binding of Y41 but not Y30. **a)** Free energy landscapes for residue Y41 sulphated (blue) and non-sulphated (red) relative to the *χ*_1_ angle of the tyrosine, showing that sulphation favours a single binding position. **b)** Representative images from across the simulation, showing the degree of motion of Y41 in its sulphated (left) and non-sulphated (right) forms. In each case, PvDBP is yellow and DARC is pink, with sulphate atoms yellow, oxygen atoms red and nitrogen atoms blue. **c)** Free energy landscapes for residue Y30 sulphated (blue) and non-sulphated (red) relative to the *χ*_1_ angle of the tyrosine, showing that sulphation disfavours a single binding position. **d)** Representative images from across the simulation, showing the degree of motion of Y30 in its sulphated (left) and non-sulphated (right) forms. **e)** The frequency of the number of contacts formed between PvDBP-RII and Y41 (left) and Y30 (right) in the sulphated (blue) and non-sulphated (red) forms during the simulations. **f)** The percentage of frames from across the simulation in which each residue from PvDBP forms interactions with Y41 (left) and Y30 (right) in the sulphated (blue) and non-sulphated (red) forms.

In the case of Y41, sulphation leads to an average free energy decrease of 7.63 kJ/mol with the decrease in free energy concentrated in the existing global minimum, at approximately *χ*_1_=-180*°*, where we observed a local decrease of 17.87 kJ/mol (Figure 2a). This corresponds with Y41 adopting a more preferred location when sulphated (Figure 2b). In contrast, while sulphation of Y30 leads to an average free energy decrease of 8.75 kJ/mol, this changes the shape of the free energy surface, with the free energy barrier at approximately *χ*_1_=-120*°* disappearing (Figure 2c) and Y30 being able to explore a wider set of *χ*_1_ configurations when sulphated (Figure 2d).

We next assessed the number of interactions formed, and the fraction of the simulation during which each interaction occurs. Using a heavy atom distance cut-off of 4.5Å to define contacts, we observe that both Y30 and Y41 make more contacts with PvDBP when sulphated. This is much more pronounced for Y41 where the average number of contacts increases from 3.64 to 7.46, whereas in Y30 we observe a shift from 1.30 to 1.87 (Figure 2e). We observe a similar pattern when considering the durability of specific interactions, with Y41 making more interactions with PvDBP than Y30 and with sulphation increasing both the number and durability of interactions (Figure 2f). In particular, Y41 makes a strong interaction with R305, which it can only reach when sulphated. We therefore find that sulphation of Y41 stabilises a specific location for the side chain in which it makes strong electrostatic interactions, while sulphation of Y30 does not lead to the formation of such a favourable contact, but instead increases the strength and durability of a smaller number of transient interactions.

### Y41-S contributes more to binding affinity and erythrocyte invasion than Y30-S

The nature of the compact binding pocket for the sulphate on Y41, and the lack of a similar binding pocket for that on Y30, lead to the prediction that the sulphate on Y41 contributes more to the binding affinity than that on Y30. To test this, we generated a set of peptides containing the ordered region of DARC ectodomain (residues 19-47) and analysed their binding to PvDBP-RII by isothermal titration calorimetry (ITC) (Figure 3a, Supplementary Figure 1). A peptide in which both Y30 and Y41 are sulphated bound to PvDBP-RII with an affinity of 58nM, while a peptide in which neither sulphate is present, bound with much lower affinity of 23μM, quantifying the importance of sulphation for this interaction. We next tested peptides in which either Y30 or Y41 is sulphated. A peptide in which Y30 is sulphated, but Y41 is not, bound with an affinity of 552nM, showing that the loss of Y41 sulphation leads to a ∼9-fold reduction in affinity. In contrast, a peptide in which Y41 is sulphated but Y30 is not, bound with an affinity of 168 nM, with a ∼3.5-fold reduction in binding affinity from the fully sulphated peptide. Therefore, both sulphates impact binding affinity, with Y41 providing a greater contribution to affinity.

**Figure 3:**
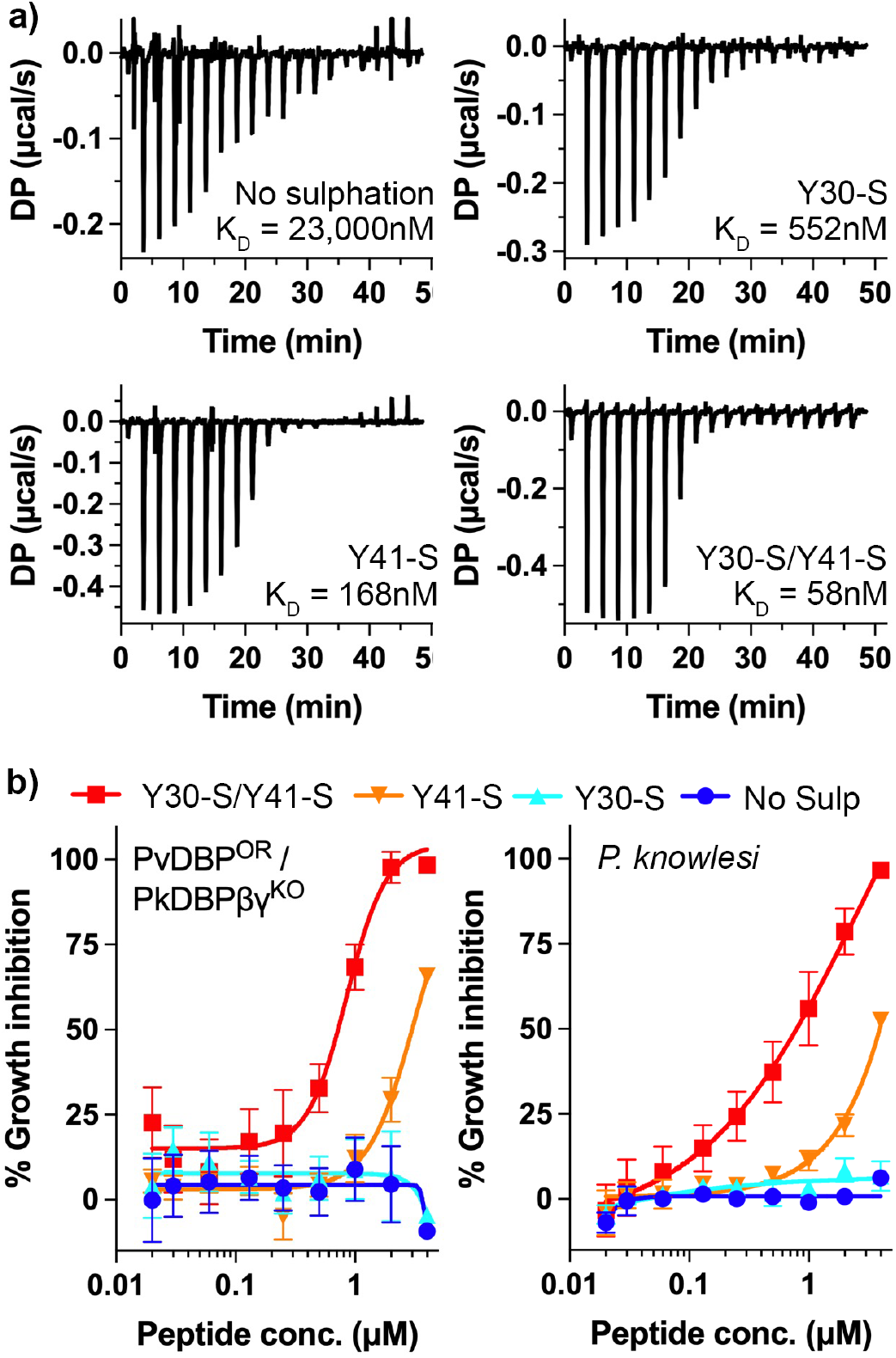
The effect of tyrosine sulphation of DARC on PvDBP-RII affinity and parasite invasion. **a)** Isothermal titration calorimetry measurements of the binding of synthetic DARC peptides to PvDBP-RII. Shown are single representative traces for non-sulphated peptide and peptides sulphated on Y30, on Y41 or on both 30 and 41. Each stated K_D_ value is the mean from n=3 technical replicates. **b)** Growth-inhibitory activity for the same four peptides in an assay which assess the growth of a *Plasmodium knowlesi* line in which PkDBPs have been replaced by PvDBP (PvDBP^OR^ / PkDBPβγ^KO^, left) or wild-type *Plasmodium knowlesi* (right). The Y30-S/Y41-S peptide inhibited with an IC_50_ of 0.72μM for PvDBP^OR^ / PkDBPβγ^KO^ and 0.79μM for *P. knowlesi*. The Y41-S peptide inhibited with an IC_50_ of 2.99μM for PvDBP^OR^ / PkDBPβγ^KO^ and 3.92μM for *P. knowlesi*. Technical replicates (n=2) from each assay were averaged, and data presented represents the mean ± standard error of the mean of four separate biological replicates (n=2 in Fy^a^ donor blood, and n=2 in Fy^b^, to account for variation between DARC alleles). IC_50_ values were identified using a variable slope four-parameter logistic curve.

We next assessed whether the differential effects of Y30 and Y41 sulphation on the affinity of DARC for PvDBP are mirrored during red blood cell invasion *in vitro*. To test this, we used a transgenic *Plasmodium knowlesi* line which has been modified to express PvDBP instead of the native, orthologous DARC binding protein, PkDBPα^13,14^. This transgenic parasite invades Duffy-positive erythrocytes in culture and is similarly affected by PvDBP-RII-targeting antibodies as *Plasmodium vivax* clinical isolates^13^. As it is currently technically impossible to generate transgenic erythrocyte lines in which DARC sulphation is specifically modulated, we instead assessed the ability of our four DARC ectodomain peptides to inhibit the growth of this transgenic PvDBP-expressing line, by measuring the effect of different concentrations of each peptide in a growth inhibition assay (Figure 3b). We found that the Y30-S/Y41-S double sulphated peptide was most effective at blocking parasite growth (IC_50_=0.72μM), followed by the Y41-S peptide (IC_50_=2.99 μM), with no growth inhibition observed for the Y30-S and non-sulphated peptides. Therefore, the effect of the peptides on parasite growth inhibition and the affinity that the peptides have for PvDBP-RII show the same pattern, with sulphation of Y41 having the greatest effect on peptide efficacy and sulphation of Y30 making a significant, but smaller difference.

We also conducted the equivalent experiment using unmodified (wild-type) *Plasmodium knowlesi* and observed a similar outcome, with IC_50_=0.79μM for the double sulphated peptide, IC_50_=3.92μM for Y41-S and no inhibition observed for non-sulphated and Y30-S peptides, highlighting the parallel in DARC binding between PvDBP and PkDBP*α*. Therefore, while a panel of PvDBP RII-targeting antibodies which blocked invasion of *P. vivax* clinical isolates and PvDBP-expressing *P. knowlesi* (including DB1 and DB9) were not cross-reactive against wild-type *P. knowlesi*^13^, the similar effect of DARC-based peptides on these two parasite lines indicates that PvDBP and PkDBP*α* engage DARC in a similar way.

### Mapping the epitope for growth-neutralising antibody DB1

We next analysed the location of the epitope for DB1 (Figure 4). This antibody inhibits growth of transgenic *Plasmodium knowlesi* parasites expressing PvDBP and of a subset of patient isolates of *Plasmodium vivax* which express different PvDBP variants^13^. Notably, DB1 binds to a site on PvDBP-RII which contains the polymorphic ^339^DEK^341^ motif^33^, making direct contact with these residues (Supplementary Figure 2), explaining in part why it is not a strain-transcendent neutralising antibody^13^. DB1 also binds to a site which does not overlap the DARC binding surface, explaining why it does not directly block DARC_ecto_ binding^13^ (Figure 4a).

**Figure 4:**
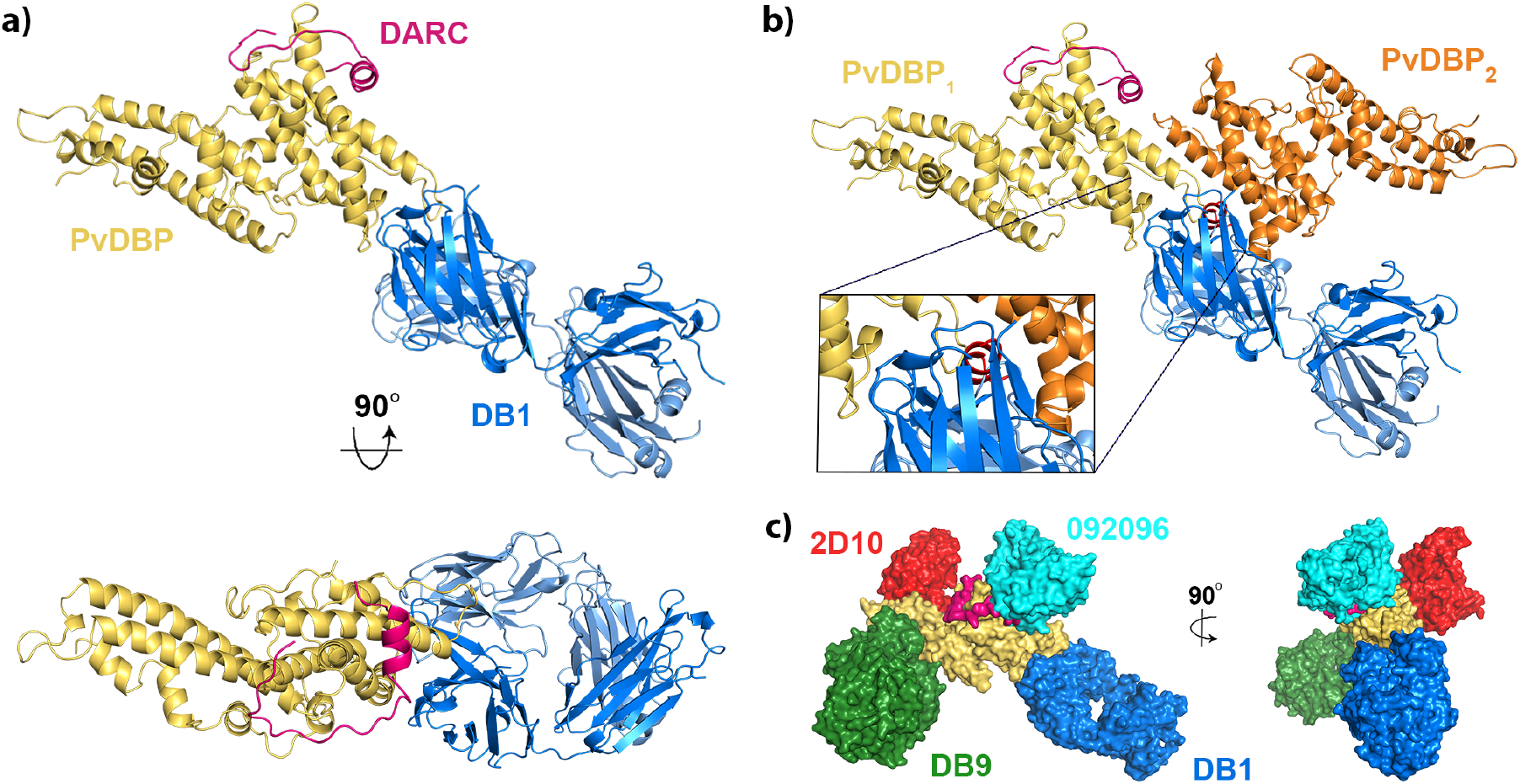
Structural basis for neutralising antibody binding to PvDBP. **a)** Structure of PvDBP-RII (yellow) bound to DARC (pink) and antibody DB1. **b)** A model of the putative PvDBP dimer (yellow and orange) bound to DARC and DB1, showing that DB1 clashes with the putative dimerization interface. **c)** A composite structure in which four different neutralising antibodies, DB1 (blue), DB9 (green)^13^, 2D10 (red)^30^ and 092096 (cyan)^20^ are docked on to the structure of PvDBP-RII (yellow) and DARC (pink).

Previous structural studies of PvDBP-RII have observed dimers to form within crystals and have also shown DARC-dependent dimerization at high concentrations in solution in the presence of a non-sulphated DARC peptide^25,29^. DB1 would block this proposed dimerization interface, with the Fab fragment overlapping with the second PvDBP copy and associated DARC peptide from the dimer (Figure 4b). To determine whether the mechanism of action of DB1 is inhibition of PvDBP-RII dimerization, we used small-angle X-ray scattering to assess the solution mass of a complex consisting of PvDBP-RII bound to Fab fragment of DB1. As predicted, the PvDBP-RII:DB1 complex showed the mass and maximum dimensions as expected for a monomeric complex (Supplementary Figure 3). We also conducted a SAXS experiment for the PvDBP-RII:DARC_19-47_ complex, but found that aggregation confounded collection of interpretable data. We therefore collected SAXS data for PvDBP-RII:DARC_19-47_ bound to Fab fragment of antibody DB9, which binds distant from the dimerization interface and would not prevent dimer formation. Both in SEC-SAXS experiments, in which the complex was passed through a size-exclusion column immediately before SAXS data collection (Supplementary Figure 3), or when studied at high concentrations of up to 6mg/ml in solution (Supplementary Figure 4) we did not observe dimerization of the PvDBP-RII:DB9:DARC_19-47_ complex. As PvDBP-RII bound to sulphated DARC_19-47_ does not dimerise in our hands, we cannot therefore conclude that DB1 acts by blocking dimerization.

## Discussion

Sulphation of DARC has been known for decades to play an important role in its binding to both cytokines and to PvDBP. However, previous structural studies used bacterially expressed DARC ectodomains, which lack tyrosine sulphation. To test whether these sulphates impact PvDBP binding, we therefore determined a structure of PvDBP-RII in complex with a sulphated DARC ectodomain. In the presence of tyrosine sulphation, we see an additional 17 residues of DARC, binding in an extended conformation around a protrusion on the PvDBP surface. We found that the sulphate on Y41 docks into a compact pocket on PvDBP, making direct interactions which are likely to be required for residues 31-47 of DARC to become ordered. Indeed, specific removal of the sulphate on Y41 leads to a ∼9-fold reduction in affinity and negates growth-inhibitory activity at the concentrations tested. In contrast, Y30 plays a less significant role in the interaction, not binding to PvDBP in the structure, dynamic in molecular dynamics simulations and with specific removal of the sulphate leading to ∼3.5- and ∼4-fold reductions in affinity and growth inhibitory activity.

Our study also reveals the epitope for the growth-neutralising antibody DB1 and shows that it binds to a polymorphic site distant from the DARC binding site, as predicted based on previous studies which show that DB1 does not block DARC binding^13^. Instead, DB1 binds in a location which would overlap with a putative dimerization interface on PvDBP-RII^25^. The growth-neutralising antibodies which target PvDBP-RII, and have known structures, bind to a variety of locations, suggesting a range of possible molecular mechanisms for neutralisation (Figure 4c). DB9^13^ and 2D10^30^ lie on subdomain 3, distant from both DARC binding site and the putative dimerization interface, 092096^20^ blocks both DARC binding and occludes the putative dimerization interface and DB1 blocks only the putative dimerization interface. However, our SAXS data revealed monomers of PvDBP-RII complexes in solution, even at high concentrations, leading us to question whether PvDBP-RII dimerization is physiological and whether blocking of dimerization is a mechanism of inhibition (Figure S2 and S3). Studies in which full-length PvDBP is assessed in the context of parasite invasion will be required to resolve whether dimerization occurs *in vivo*.

Our study, therefore presents a complete view of the interaction between DARC and PvDBP-RII and reveals the role of DARC sulphation on PvDBP binding. However, our view of the complete PvDBP protein remains incomplete. Studies of PvDBP have long focused on the RII domain, through ease of production and due to the demonstration that this domain contains the DARC binding site. However, questions remain, which will only be answered when we understand the structure of full-length PvDBP and how it binds to full-length DARC. For example, it is also possible that the putative dimerisation interface observed in PvDBP-RII has other roles in full-length PvDBP, which are affected by antibody binding. Further study is also required to understand the role of the DARC Fy^a^/Fy^b^ polymorphism on PvDBP binding as the role of the polymorphic residue 42 is not certain from this study. Nevertheless, we now reveal, for the first time, the full interaction between the ectodomain of DARC and PvDBP-RII, showing the details of this interaction critical for red blood cell invasion in *Plasmodium vivax*. This identifies new surfaces of PvDBP which could be targeted by small molecules or antibodies that block the interaction with DARC, or that can be included in vaccine immunogens. These findings will guide future design of therapeutic agents to target the scourge of malaria.

## Materials and Methods

### Protein expression and purification

For crystallisation, the coding sequence for PvDBP-RII (D194-S508) was cloned into the pHLsec vector, which included a C-terminal 6xHis tag. DARC ectodomain (M1-S60, numbered as for the mature polypeptide) was cloned into the pENTR4 vector containing a C-terminal TEV protease cleavage site and 6xHis tag. DB1 heavy and light chains were supplied in the AbVec-hIgG1 and AbVec-hIgKappa vectors respectively. The constructs were transfected into Expi293 cells (Thermo Fisher Scientific) using the ExpiFectamin 293 Transfection Kit (Gibco). The transfected cells were collected after 4 days and the supernatant clarified by centrifugation at 10,000g and supplemented with 150 mM NaCl and 20 mM imidazole (final concentration).

For isothermal titration calorimetry, PvDBP-RII (D194-T521) was expressed with an N-terminal monomeric Fc domain (mono-Fc). Three alanine substitution mutations, T257A, S353A and T422A, were introduced to the PvDBP-RII sequence to remove potential N-linked glycosylation sites. The construct was transfected into Expi293 cells as above and harvested after 5 days. The mono-Fc tagged PvDBP-RII was purified using HiTrap Protein A HP column (Cytiva).

DB1 antibody was purified by binding clarified cell supernatant to Protein A Agarose beads (Thermo Fisher Scientific), with elution into 100 mM glycine pH 3.0 and was quickly neutralised with 1 M Tris-HCl pH 8.0. To generate DB1 Fab fragments, purified DB1 was buffer exchanged into 150 mM NaCl, 20 mM HEPES pH 7.5, 20 mM L-cysteine for overnight cleavage at 37°C with immobilized papain, followed by protein-A agarose purification to remove Fc and uncleaved DB1 antibody.

PvDBP-RII and DARC ectodomain used for crystallisation was purified by immobilized-metal-affinity chromatography by batch binding clarified cell supernatant to Ni-NTA resin, with elution into 300 mM imidazole, 150mM NaCl, 20mM HEPES pH 8.0 and was buffer exchanged into 150 mM NaCl, 20 mM HEPES pH 7.5 using a Vivaspin 3kDa column. TEV protease and PNGase F were added to DARC ectodomain overnight at 4°C to remove the TEV-6xHis tag and glycans, respectively.

### Crystallisation, data collection, and structure determination

Crystallisation trials were carried out using vapour diffusion in sitting drops, mixing 100 nL protein complex solution and 100 nL well solution. Crystals were visible from day 12 with a well solution of 15% w/v jeffamine D-2003 and 10% v/v ethanol. The crystals were cryoprotected by transfer into drops of well solution supplemented with 25% glycerol, and were cryo-cooled in liquid nitrogen for data collection.

Data was collected at the I24 beamline at Diamond at a wavelength of 0.9999 Å. After processing by autoPROC, the dataset was scaled using AIMLESS (v.0.73)^34^, producing a complete dataset at a resolution of 2.5 Å. The structure was solved by molecular replacement with Phaser MR (v.2.8.3)^35^ using a search model for PvDBP-RII (PDB: 6R2S). The model was built and refined using cycles of COOT (v.0.8.9.2)^36^ and BUSTER (v.2.10)^37^.

### Molecular dynamics simulations

All simulations performed using OpenMM v7.7^38^. Models of PvDBP (residues 217-509) and DARC (residues 19-47) were capped at C- and N-termini using N-methyl and acetyl groups, respectively, protonated at pH of 7.5 using H++ ^39^ and were soaked in truncated octahedral water boxes with a padding distance of 1 nm. NaCl was added to 150 mM while neutralising charges as described in^40^. Systems were parameterised using the Amber14-SB force field^41^ and water modelled using TIP3P-FB^42^ and tleap^43^. For sulphotyrosine residues, we used parameters from ^44^. Three systems were prepared from our crystal structure, with both Y30 and Y41 sulphated, with only Y30 sulphated and with only Y41 sulphated.

Non-bonded interactions were calculated using the particle mesh Ewald method^45^ with a real-space cut-off of 0.9 nm and error tolerance of 0.0005. Water molecules and heavy atom-hydrogen bonds were rigidified using SETTLE^46^ and SHAKE^47^ algorithms, respectively. Hydrogen mass repartitioning^48^ was used to allow 4 fs time steps. Simulations were run using the mixed-precision CUDA platform in OpenMM using the Middle Langevin Integrator with a friction coefficient of 1 ps^-1^ and the Monte-Carlo Barostat at a pressure of 1 atm. We equilibrated systems using a multi-step protocol: (i) energy minimisation over 10,000 steps, (ii) heating of the NVT ensembles from 100K to 300K over 200 ps, (iii) 200 ps simulation of the NPT ensembles at 300K, (iii) cooling of the NVT ensembles from 300K to 100K over 200 ps, (iv) energy minimisation over 10,000 steps, (v) heating of the NVT ensembles from 100K to 300K over 200ps, and (vi) 5 ns simulation of the NPT ensembles at 300K.

We initialised well-tempered metadynamics using Plumed^49^. The first pair of simulations was biased on the *χ*_1_ and *χ*_2_ angles of DARC position 30. These were initialised with structures with both Y30 and Y41 sulphated and with just Y41 sulphated, using an initial hill height of 1.2 kJ/mol, a bias factor of 10, and bias widths of 0.07 rad, with biases being deposited every 500 steps. The second pair of simulations was biased on the *χ*_1_ and *χ*_2_ angles of DARC position 41. These were initialised with structures with both Y30 and Y41 sulphated and with just Y30 sulphated, using an initial hill height of 3.0 kJ/mol, a bias factor of 10, and bias widths of 0.07 rad, with biases being deposited every 500 steps. We found that a lower hill height of 1.2 kJ/mol was insufficient to overcome free energy barriers in the system. For the second pair of simulations, we applied harmonic restraints on the backbone positions of the DARC peptide using the crystal structure as the reference and a force constant of 1500 kJ/(mol nm^2^). Production simulations were performed for 500 ns and convergence was assessed using block error analysis on the computed free energy surfaces with respect to *χ*_1_ and *χ*_2_ angles of the biased positions. To recover unbiased ensembles from our simulations, we calculated weights (*w*) for each configuration (*s*) using the time-independent reweighting scheme^50^

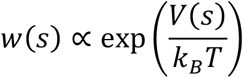

where *V*(*s*) is the configuration-dependent bias at the end of the simulation and *k*_*B*_*T* is the product of the Boltzmann constant and temperature. We used MDTraj (v. 1.9.6)^51^ to analyse trajectories.

### Isothermal titration calorimetry

Isothermal titration calorimetry (ITC) experiments were performed on a MicroCal PEAQ-ITC (Malvern). PvDBP-RII used in the ITC experiments was tagged N-terminally with a monomeric Fc domain (mono-Fc) and contained T257A, S353A and T422A substitutions to remove N-linked glycosylation sites; modifications are not in close proximity to the DARC binding site and hence should not interfere with binding. All synthetic peptides used in the ITC experiments were purchased from Cambridge Research Biochemicals (Billingham, UK) with the purity level of 95%. Prior to the measurement, mono-Fc tagged PvDBP-RII was extensively dialyzed against 1x phosphate-buffered saline (PBS), and the peptides were dissolved in the same dialysis buffer to prevent buffer mismatch. The peptides were placed in the syringe at a concentration of ∼130 μM for Ac-QLDFEDVWNSS-[Y(SO3H)]-GVNDSFPDGD-[Y(SO3H)]-DANLEL-NH_2_, ∼300 µM for Ac-QLDFEDVWNSSYGVNDSFPDGDYDANLEL-NH2, ∼200 µM for Ac-QLDFEDVWNSS-[Y(SO3H)]-GVNDSFPDGDYDANLEL-NH_2_, and ∼260 µM for Ac-QLDFEDVWNSSYGVNDSFPDGD-[Y(SO3H)]-DANLEL-NH_2_, whereas the concentration of mono-Fc tagged PvDBP-RII in the cell was ∼20 μM for all experiments. The peptide concentrations were determined by BCA assay using the Pierce Rapid Gold BCA Protein Assay Kit (ThermoFisher Scientific), and the protein concentration was determined by UV absorbance at 280 nm. The titrations were all performed with peptides in the syringe and mono-Fc tagged PvDBP-RII in the cell and consisted of a single 0.4 µL injection followed by 18 injections of 2 μL with injection duration of 4 s, injection spacing of 150 s, stir speed of 750 rpm, and reference power of 10 μcal/s. Experiments were conducted at 25°C in triplicate (n = 3), and the data are reported as the arithmetic mean ± SD. Fitting of the integrated titration peaks was performed with MicroCal PEAQ-ITC Analysis Software (Malvern) provided with the instrument, then all data were exported to and plotted in Prism 9 (Dotmatics).

### Small angle X-ray scattering

Mono-Fc tagged PvDBP-RII used for SAXS experiments was expressed in Expi293 cells (Thermo Scientific) as described above and purified using the CaptureSelect C-tagXL Affinity Matrix (Thermo Scientific). The elution was concentrated and buffer-exchanged into 20 mM HEPES, 150 mM NaCl pH 7.5 using a PD-10 Desalting Column (Cytiva) before TEV cleavage (1:100 TEV to protein ratio) overnight at 4*°*C. After cleavage, mono-Fc was removed using Pierce Protein A agarose beads (Thermo Scientific). The flow-through containing PvDBP-RII was then mixed with DB1 or DB9 Fab (1.5:1 Fab to protein ratio) for 1 hr before concentration and size-exclusion chromatography on a Superdex 200 Increase 10/300 GL column (Cytiva). The fractions containing PvDBP-RII with Fabs were concentrated or were mixed with DARC_19-47_ Y30-S/Y41-S double sulphated peptides in case of the PvDBP-RII-DB9 Fab complex for 1 hr prior to concentration.

Small-angle X-ray scattering (SAXS) data were collected at beamline B21 at the Diamond Light Source (Didcot, UK). Scattering was detected using X-rays at a wavelength of 0.99 Å and an Eiger 4M detector with a detector-sample distance of 4.014 m. For the PvDBP-RII:DB9 Fab complexes with or without Y30-S/Y41-S double sulphated peptides, experiments were performed via two methods: 1) batch mode where 35 µL of samples at 6, 3, 1 mg/mL concentrations with corresponding matching buffers were injected directly into the capillary, using the EMBL Arinax sample handling robot, for data collection, and 2) SEC-SAXS where 50 µL of 6 mg/mL samples were gel filtered on the Shodex PROTEIN KW403-4F column (Shodex Denko Europe) equilibrated with 20 mM HEPES, 150 mM NaCl, pH 7.5 prior to data collection. The data for PvDBP-RII:DB1 complex were only collected using the SEC-SAXS method at the starting concentration of 6 mg/mL.

The data were processed using ScÅtter^52^ with the ATSAS software suites^53^. For SEC-SAXS, buffer frames were averaged and subtracted from averaged frames of the complex peak fractions. The radius of gyration (Rg), distance distribution function P(r) and the maximum particle diameter (Dmax) were determined using ScÅtter. Twenty-nine *ab initio* initial bead models were generated with DAMMIF^53^ from the ATSAS software suites in ScÅtter and averaged with DAMAVER^54^ followed by refinement against the original data using DAMMIN^55^. The 20 Å envelope of the refined bead model was generated in Chimera^56^. Crystal structures of PvDBP-RII in complex with DB9 Fab^13^ (PDB: 6R2S) overlayed with the Y41-S DARC peptide structure obtained in this study and the crystal structure of PvDBP-RII with DB1 Fab from the current study were fitted into the envelopes using Chimera.

CRYSOL^53^ was used to derive theoretical scattering data from monomeric and dimeric version of PvDBP-RII:DB9 or PvDBP-RII:DB1 complexes and to compare these data with the experimental results.

### Growth inhibition assays with transgenic Plasmodium knowlesi

To investigate DBP-DARC interactions *in vitro*, the variably sulphated DARC ectodomain peptides described above were used to competitively inhibit the host-parasite interaction between erythrocytic DARC and parasite-expressed DBP, which is essential for invasion and subsequent replication. Growth inhibition assays were run as previously described in^13,14^ with parasite growth measured via lactate dehydrogenase (LDH) assay after one intraerythrocytic cycle (approx. 27h) in the presence of various peptides, prepared in 2-fold dilution curves. Each 96-well plate included internal controls (infected, untreated erythrocytes and uninfected erythrocytes) which were used to calculate the percentage growth inhibition in each treatment well using the following formula^13,14^ –

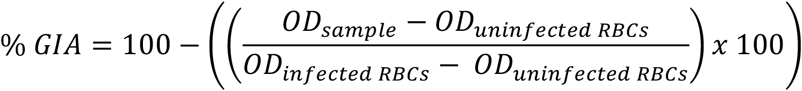

Assays were conducted with a minimum of 2 technical replicates, and data presented represents averages from four separate experiments (n=2 each using Fy^a^ and Fy^b^ donor red blood cells). Assays were performed using both *P. knowlesi* wild-type A1-H.1 (expressing PkDBPα), as well as transgenic *P. knowlesi* PvDBP^OR^ / PkDBPβγ^KO^, which expresses the *P. vivax* DBP as an orthologue replacement of the homologous PkDBPα. IC_50_ values were identified using a variable slope four-parameter logistic curve, calculated using GraphPad Prism v9.4.1.

## Acknowledgements

MKH is a Wellcome Trust Investigator (220797/Z/20/Z). RM and TP are funded by the Skaggs-Oxford graduate scholarship. B.G. is funded by the Wellcome PhD program in Cellular Structural Biology. S.M.D. is supported by a UK Medical Research Council LID PhD studentship. R.W.M was supported by the UK Medical Research Council (MRC Career Development Award MR/M021157/1). The authors thank Simon Draper for the coding sequence for antibody DB1 and the plasmid construct for monoFc-tagged PvDBP-RII. We thank the staff at Diamond beamline I04 for help with crystallographic data collection and staff at Diamond beamline B21 for help with SAXS data collection.

## Author contributions

R.M. produced proteins and R.M., E.L. and M.K.H. determined the crystal structure. T.P. conducted ITC and SAXS experiments. B.G. performed molecular dynamics analysis. S.M.D., F.M. and R.W.M. conducted parasite growth-inhibition experiments. R.W.M. and M.K.H. devised the study. M.K.H. drafted the manuscript and all authors discussed the results and commented on the manuscript.

## Data Availability

Coordinates and structure factors are deposited in the Protein Data Bank with accession code 8A44 and all other data and protein constructs are available from the authors on request.

---

**Supplementary Table 1:** Crystallographic statistics.

|  |  |
| --- | --- |
| **Data collection** |  |
| Space group | P2<sub>1</sub>2<sub>1</sub>2<sub>1</sub> |
| Cell dimensions: |  |
| a, b, c (Å) | 52.20, 97.95, 287.99 |
| $\alpha$, $\beta$, $\gamma$ (°) | 90, 90, 90 |
| Resolution (Å) | 92.73 – 2.49 (2.57 – 2.49) |
| Total observations | 99952 (8634) |
| Total unique | 52982 (4488) |
| R<sub>pim</sub> | 0.070 (0.735) |
| CC<sub>1/2</sub> | 0.993 (0.463) |
| I/$\sigma$(I) | 5.7 (0.7) |
| Completeness (%) | 100 (100) |
| Multiplicity | 1.9 (1.9) |
| Wilson B factor (Å<sup>2</sup>) | 56.4 |
| **Refinement** |  |
| Number of reflections | 52884 |
| R<sub>work</sub> / R<sub>free</sub> | 0.198 / 0.225 |
| Average B factor (Å<sup>2</sup>) | 67.0 |
| Number of residues: |  |
| Amino acid residues | 760 |
| Waters | 250 |
| RMSZ deviations |  |
| Bond lengths | 0.008 |
| Bond angles | 0.985 |
| Ramachandran plot |  |
| Favoured (%) | 96.3 |
| Allowed (%) | 3.7 |
| Outliers (%) | 0 |

**Supplementary Figure 1:**
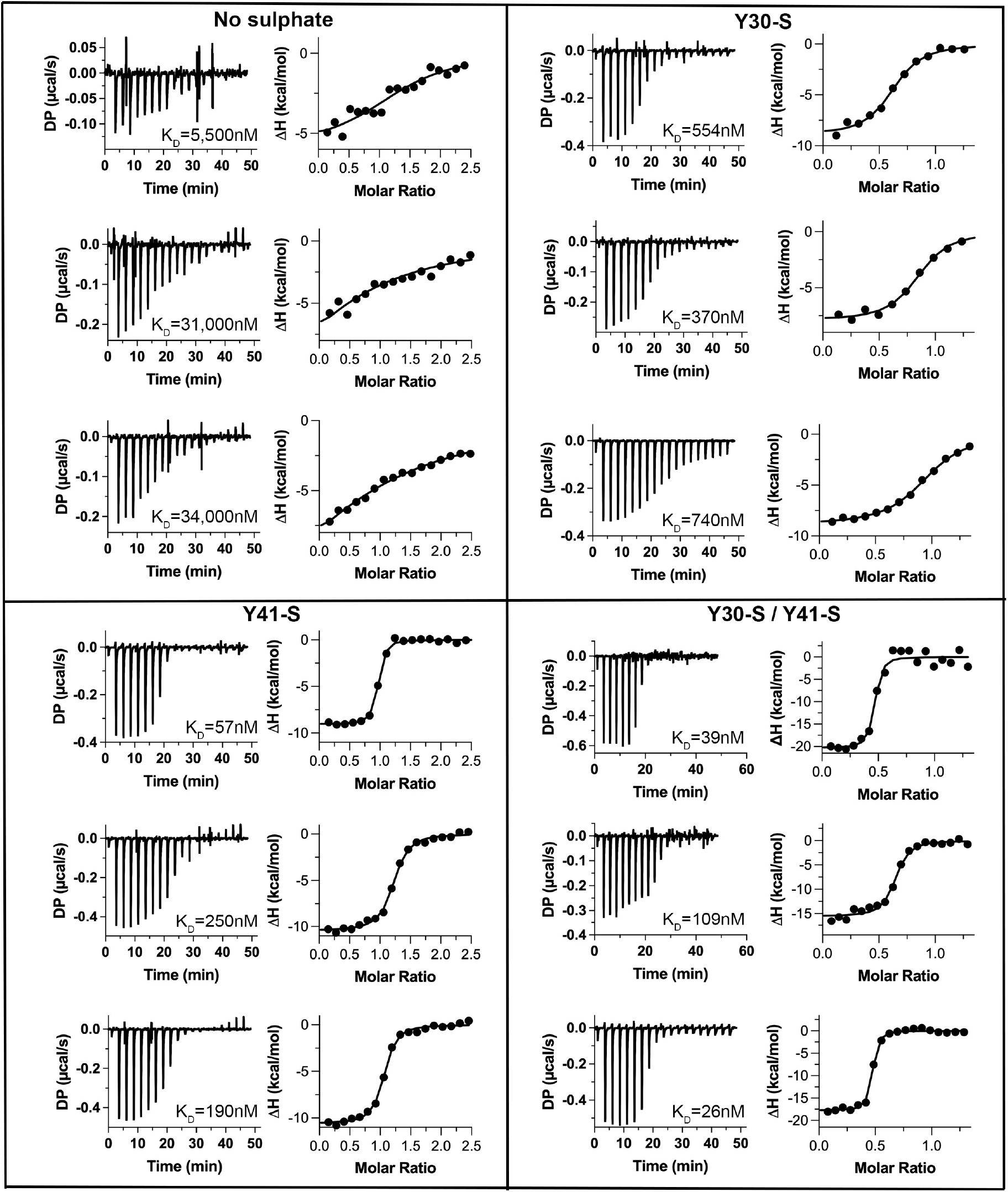
Measurement of affinities of PvDBP-RII for DARC peptides by isothermal titration calorimetry. In each case, three separate technical replicates were performed and the mean and standard deviation K_D_ value was determined. For the non-sulphated peptide, the measured K_D_ values were 5.5±6.4μM, 31±28μM and 34±40μM with the mean of 23±16μM. For the Y30-S peptide, the measured K_D_ values were 544±116nM, 370±120nM and 740±84nM with the mean of 552±186nM. For the Y41-S peptide, the measured K_D_ values were 57±13nM, 250±45nM and 190±34nM with the mean of 168±101nM. For the Y30-S / Y41-S peptide, the measured K_D_ values were 39±25nM, 109±33nM and 26±7nM with the mean of 58±45nM.

**Supplementary Figure 2:**
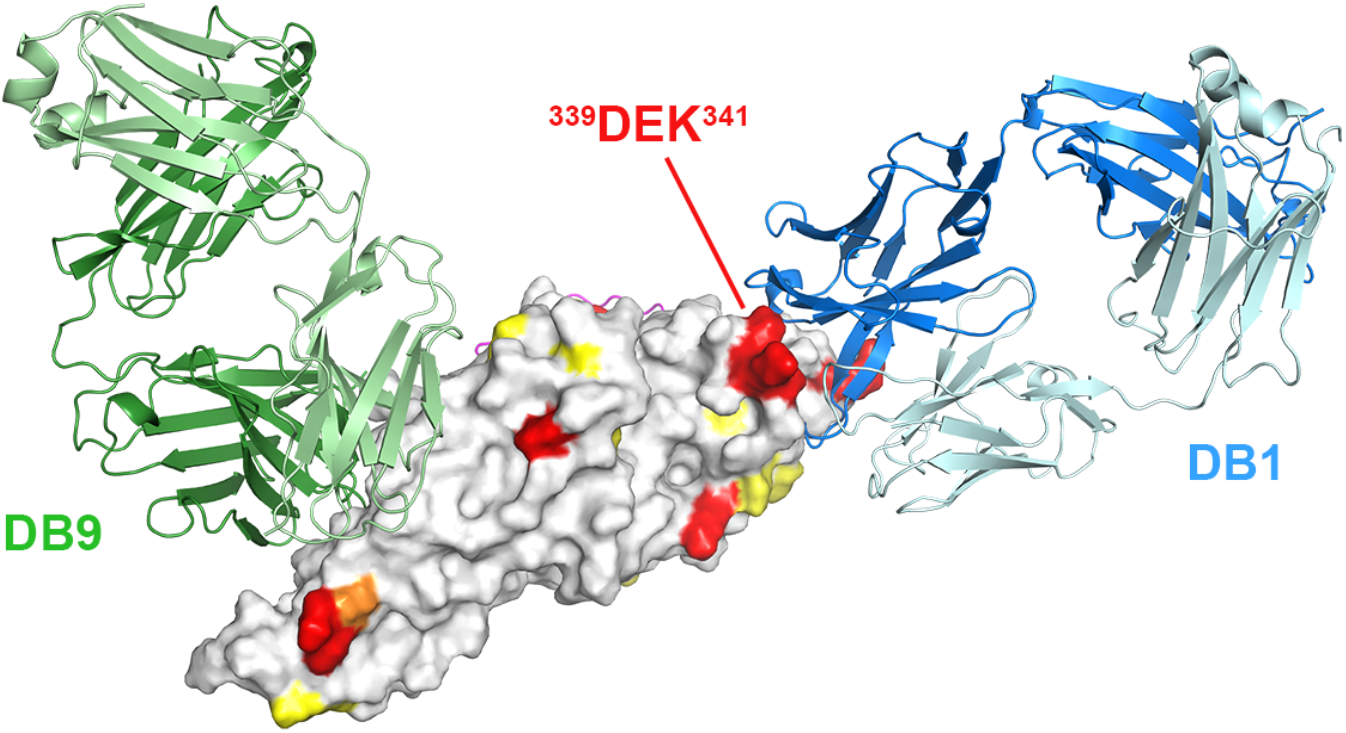
Polymorphism in the epitopes for DB1 and DB9. The structure of PvDBP-RII is shown in surface representation, which has been coloured according to sequence polymorphism (grey <0.15; yellow = 0.15-0.30; orange = 0.30-0.40 and red > 0.4). The Fab fragments of DB9 (green) and DB1 (blue) are shown. DB9 binds to a non-polymorphic site, while DB1 binds directly to the polymorphic ^339^DEK^341^ motif.

**Supplementary Figure 3:**
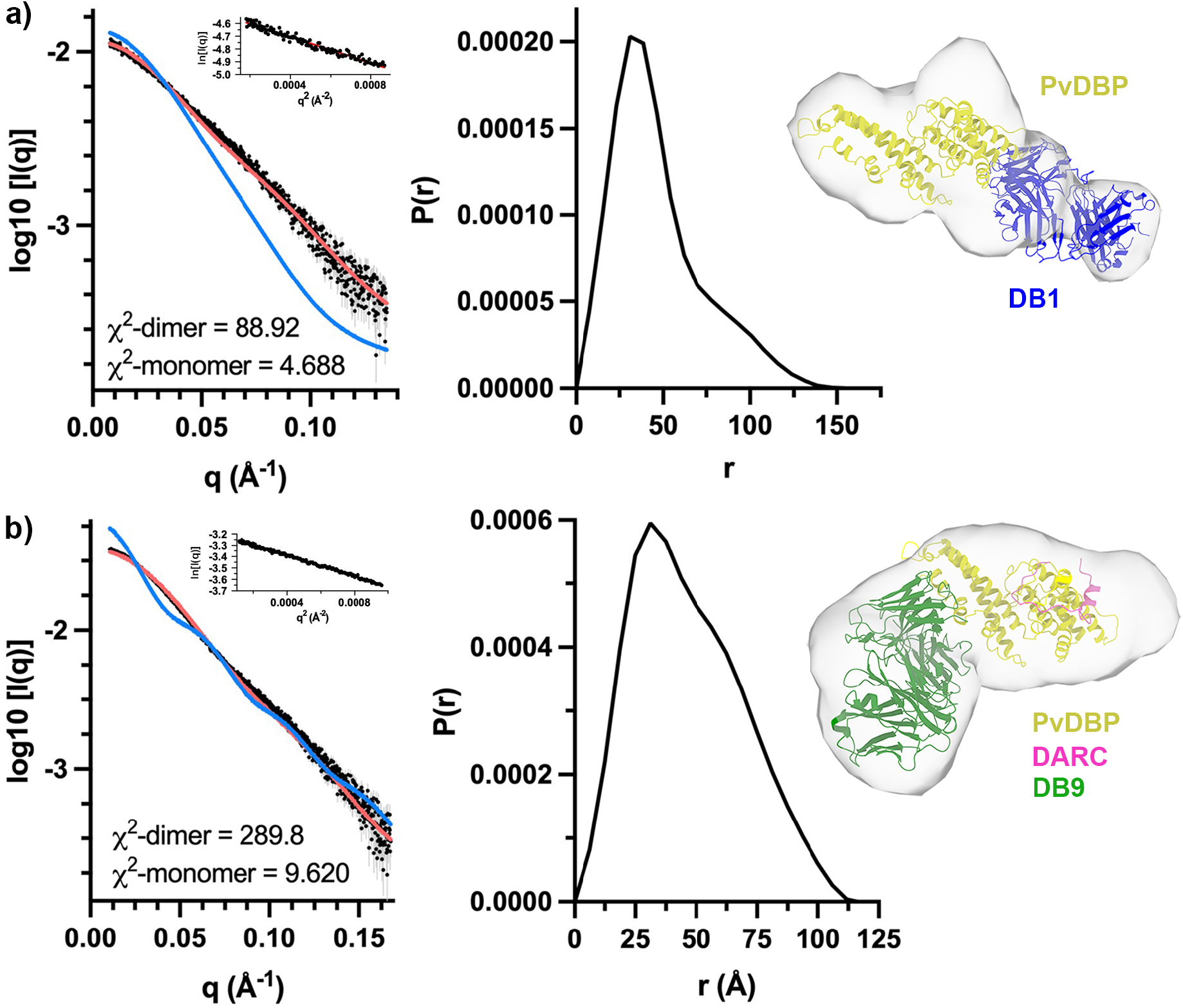
Small angle X-ray scattering data (SEC-SAXS) for PvDBP-RII complexes. Small angle X-ray scattering data (SEC-SAXS) for **a)** the complex of PvDBP-RII and the Fab fragment of antibody DB1 and **b)** the complex of PvDBP-RII, DARC_19-47_ and the Fab fragment of antibody DB9. In each case, the left-hand panel shows the refined scattering data (black circles) fitted either with the predicted scattering for the monomeric (red) or dimeric (blue) complex. The inset is the Guinier plot. The central panel shows the pair distance distribution function while the right-hand panel shows the calculated envelope into which has been docked the monomeric model.

**Supplementary Figure 4:**
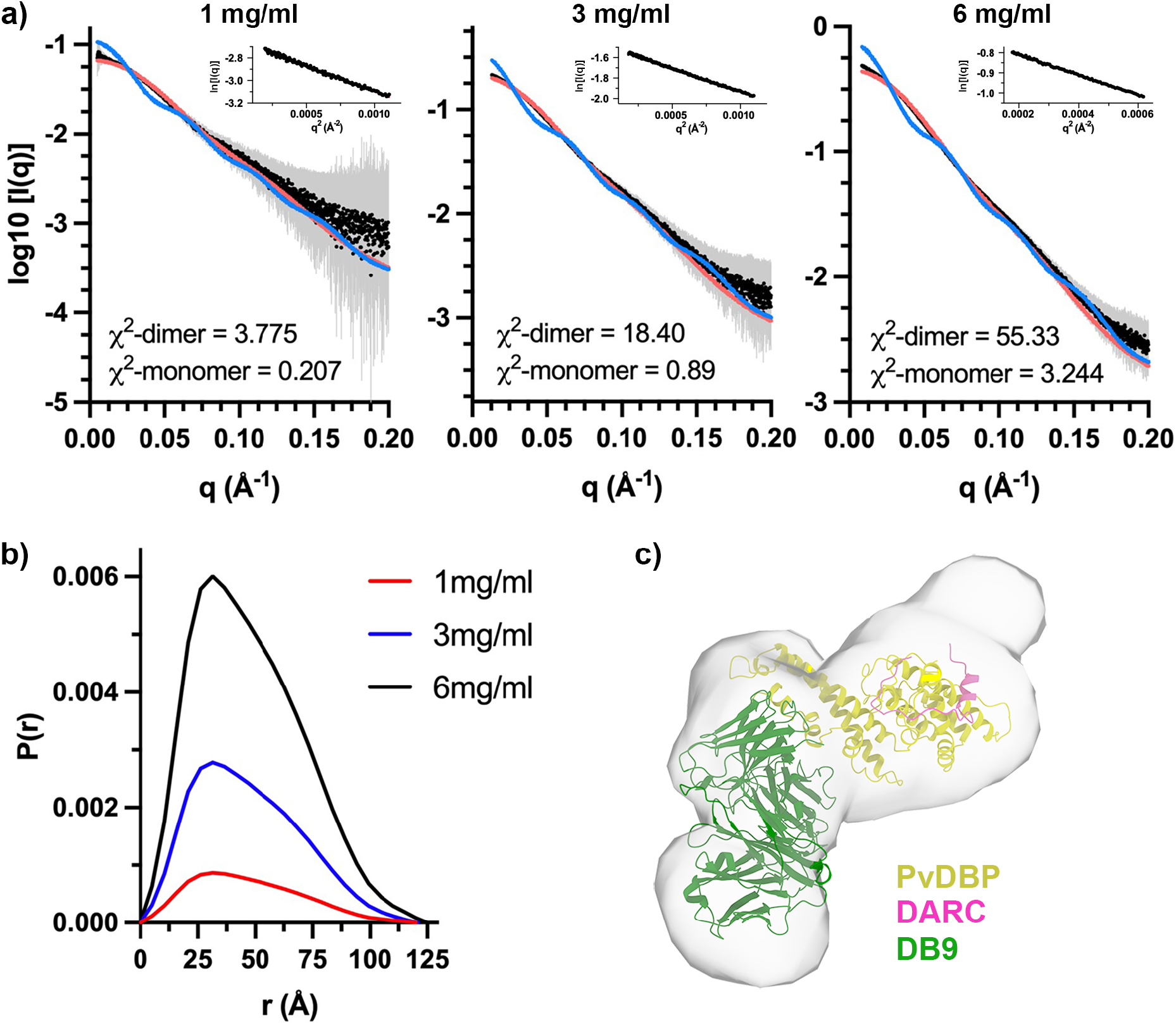
Small angle X-ray scattering data for PvDBP-RII:DARC_19-47_:DB9 complexes in batch model. **a)** Small angle X-ray scattering data (SEC-SAXS) for the complex of PvDBP-RII, DARC_19-47_ and the Fab fragment of antibody DB9 collected in batch mode at 1mg/ml (left), 3mg/ml (central) and 6mg/ml (right). In each case, the refined scattering data (black circles) is shown, fitted either with the predicted scattering for the monomeric (red) or dimeric (blue) complex. The insets are the representative Guinier plots. **b)** The pair distance distribution functions for the 1mg/ml (red), 3mg/ml (blue) and 6mg/ml (black) data. **c)** The calculated envelope into which has been docked the monomeric model from the 3mg/ml data.

